# Enhancing Gene Set Overrepresentation Analysis with Large Language Models

**DOI:** 10.1101/2024.11.11.621189

**Authors:** Jiqing Zhu, Rebecca Y. Wang, Xiaoting Wang, Ricardo Azevedo, Alexander Moreno, Julia A. Kuhn, Zia Khan

**Affiliations:** Alector, Inc.

## Abstract

Gene set overrepresentation analysis is widely used to interpret high-throughput transcriptomics and proteomics data, but traditional methods rely on static, human-curated gene set databases that lack flexibility. We introduce *llm2geneset*, a framework that leverages large language models (LLMs) to dynamically generate gene set databases tailored to input query genes, such as differentially expressed genes (DEGs) and a biological context specified in natural language. These databases integrate with methods, such as overrepresentation analysis (ORA), to assign biological functions to input genes. Benchmarking against human-curated gene sets demonstrates that LLMs generate gene sets comparable in quality to those curated by humans. *llm2geneset* also identifies biological processes represented by input gene sets, outperforming traditional ORA and direct LLM prompting. Applying the framework to RNA-seq data from iPSC-derived microglia treated with a TREM2 agonist highlights its potential for flexible, context-aware gene set generation and improved interpretation of high-throughput biological data. *llm2geneset* is available as open source at https://github.com/Alector-BIO/llm2geneset and via a web interface at https://llm2geneset.streamlit.app.

## Introduction

Gene sets, which group genes or proteins based on shared biological functions, pathways, or regulatory mechanisms, are essential tools in transcriptomics and proteomics for interpreting complex datasets[1,2]. By focusing on coordinated biological processes rather than individual genes, gene set analysis enhances the understanding of underlying biology and helps identify patterns that may be obscured in high-dimensional data. This approach allows molecular changes, such as differentially expressed genes (DEGs), to be connected to specific biological processes, supporting investigations into normal physiology, disease mechanisms, and therapeutic strategies. The generation and interpretation of gene sets are critical for deriving meaningful insights from transcriptomics and proteomics studies and for deepening our understanding of biological systems.

Methods for gene set enrichment analysis, such as overrepresentation analysis (ORA), are widely used to evaluate whether gene sets are significantly associated with a list of DEGs [3–7]. These methods typically rely on external, human-curated gene set databases, which must be regularly updated to incorporate advances in scientific knowledge. However, static databases often pose challenges when tailoring gene sets to specific research questions or experimental contexts, and the choice of gene sets can strongly influence the interpretation of transcriptomics and proteomics studies [1]. In this paper, we demonstrate how large language models (LLMs) enable the generation of gene sets using natural language. This approach supports the customization of gene sets for specific research questions and enhances the identification of biologically relevant processes when integrated with enrichment analysis.

LLMs have recently found several uses in the analysis of transcriptomics data sets and their application to problems in computational biology is an active area of research. LLMs have been used to automate cell type annotation in single cell data sets on the basis of their ability to capture marker genes of known cell types [8]. Recent work has demonstrated that LLMs can discern functional relationships within a given set of genes, yet the statistical relevance of these LLM derived observations has not been considered [9]. LLMs have been augmented with bioinformatic specific tools to solve tasks [10]. Yet, use and evaluation of LLMs as a tool for flexible gene set generation and overrepresentation analyses has not been explored.

In this study, we introduce *llm2geneset*, a framework that leverages LLMs to generate a gene set database tailored to input query genes (e.g., DEGs) and a biological context specified in natural language (Fig. 1). This dynamically generated database can be used by gene set analysis methods, such as overrepresentation analysis (ORA), to assign biological processes and functions to the input genes. We evaluate two key aspects of the framework: its ability to accurately generate gene sets from natural language descriptions and its ability to map input gene sets to known functions. Using human-curated gene sets, we test whether LLMs can recover genes curated by humans when provided with a natural language description and find that they perform comparably to human curators, with prompting strategies tuning precision. Additionally, we evaluate whether *llm2geneset* can identify the biological processes or functions represented in input gene sets, whether corresponding to single or multiple processes, and find that it outperforms traditional ORA and direct LLM prompting. Finally, we demonstrate the framework’s potential as a flexible and steerable tool for gene set analysis by applying it to transcriptomics data from iPSC-derived microglia treated with a TREM2 agonist.

**Figure 1.**
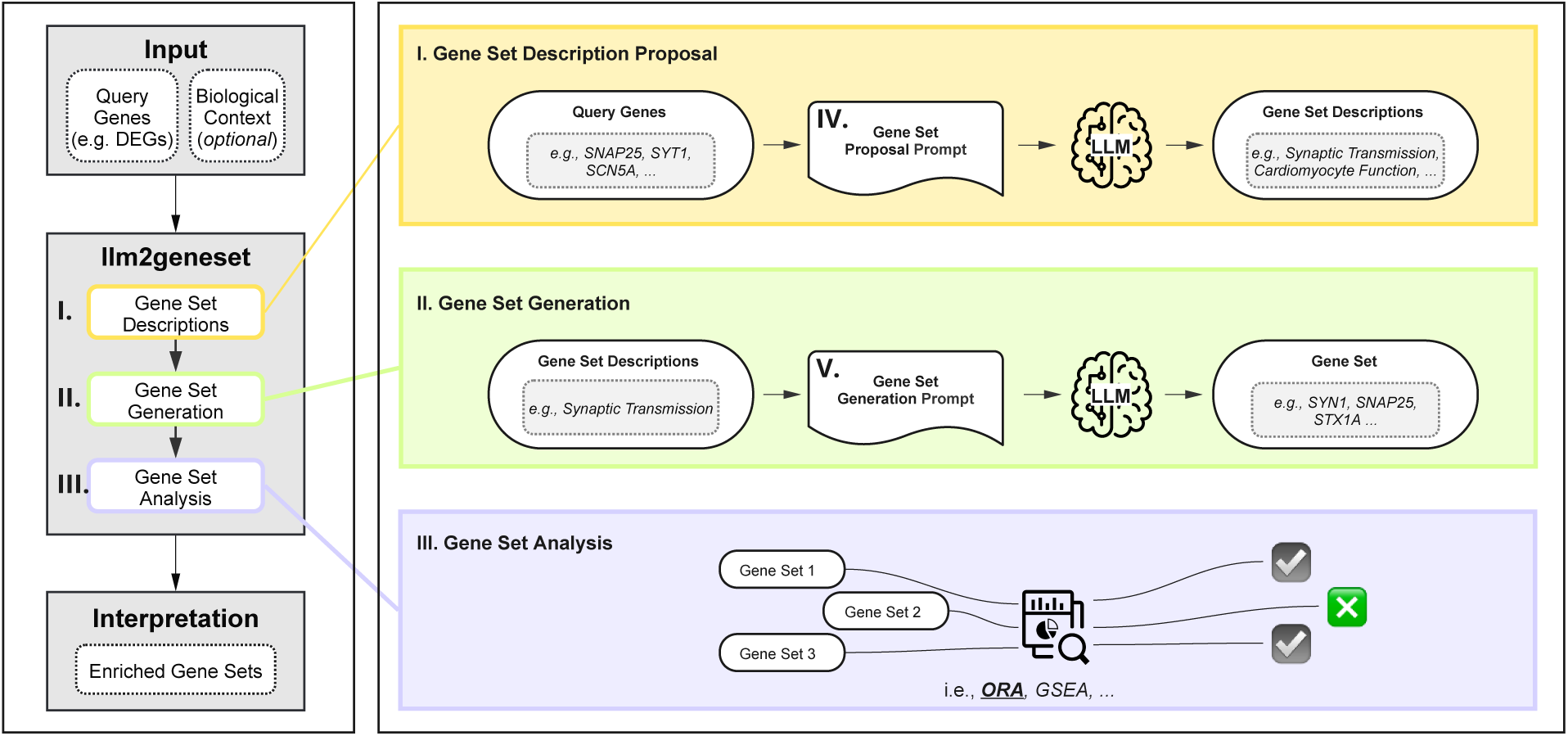
Overview of the *llm2geneset* framework. The framework takes a set of query genes (e.g., differentially expressed genes (DEGs)) and an optional biological context as input. The LLM-based gene set database construction involves two stages: first, proposing biological processes relevant to the query genes within the context, and second, generating gene sets by associating relevant genes with these processes. The resulting customized gene set database can be utilized by gene set analysis methods, such as overrepresentation analysis (ORA), to identify enriched gene sets.

## Results

### LLMs Generate Informative Gene Sets

We evaluated whether LLMs could be used to generate gene sets using flexible natural language descriptions of these gene sets. To achieve this, we used a fixed text prompt with a field that can be replaced by the natural language description of the gene set (see Methods) [11,12]. Our prompt consisted of several elements: (1) a role-prompt, a strategy whereby a system message is used to guide the generations of an LLM; (2) the prompt itself requesting generation of genes for a given biological process or cellular component; (3) a formatting component to guide the LLM to produce output using standard HUGO Gene Nomenclature Committee (HGNC) gene symbols that can be programmatically loaded into data structures for downstream use in gene set tests (see Methods).

To evaluate the output of an LLM, we assessed whether the gene sets generated by an LLM were significantly overrepresented in corresponding human curated gene sets (Fig. 2A). To compute whether the overrepresentation we observed was not due to chance, we used the hypergeometric distribution to compute a p-value for a one-tailed Fisher’s exact test and adjusted for multiple testing (see Methods). We performed this assessment using natural language descriptions of gene sets in 3 human curated gene set databases: KEGG [13], Reactome [14], WikiPathways [15], and 1000 randomly sampled gene sets from Gene Ontology Biological Process (GOBP) database [16]. Curated for different applications, each gene set database captured differing underlying biology and varied in total number of gene sets (Fig. S2A). In total, we examined overrepresentation in 3,939 gene sets and the gene sets varied in size ranging from a median of 24-78 genes (Fig. S2B). We evaluated LLM gene set generations from 3 different language models: GPT-3.5, GPT-4o-mini, and GPT-4o [17,18]. We found that between 33% to 94% LLM generated gene sets were overrepresented in human curated gene sets at a Bonferroni adjusted p-value of 0.01 (Fig. 2B). We observed a higher fraction of overrepresented gene sets generated by the more capable GPT-4o model than GPT-3.5. This fraction tracked with the relative capabilities of each model we considered with GPT-4o mini ranking between GPT-4o and GPT-3.5. Overall, all models performed worst on the GOBP gene set database. Across most gene sets, the over-representation p-value was smallest for GPT-4o indicating that on an individual gene set level a more capable model yields gene sets in greater agreement with curated gene sets (Fig. 2C).

**Figure 2.**
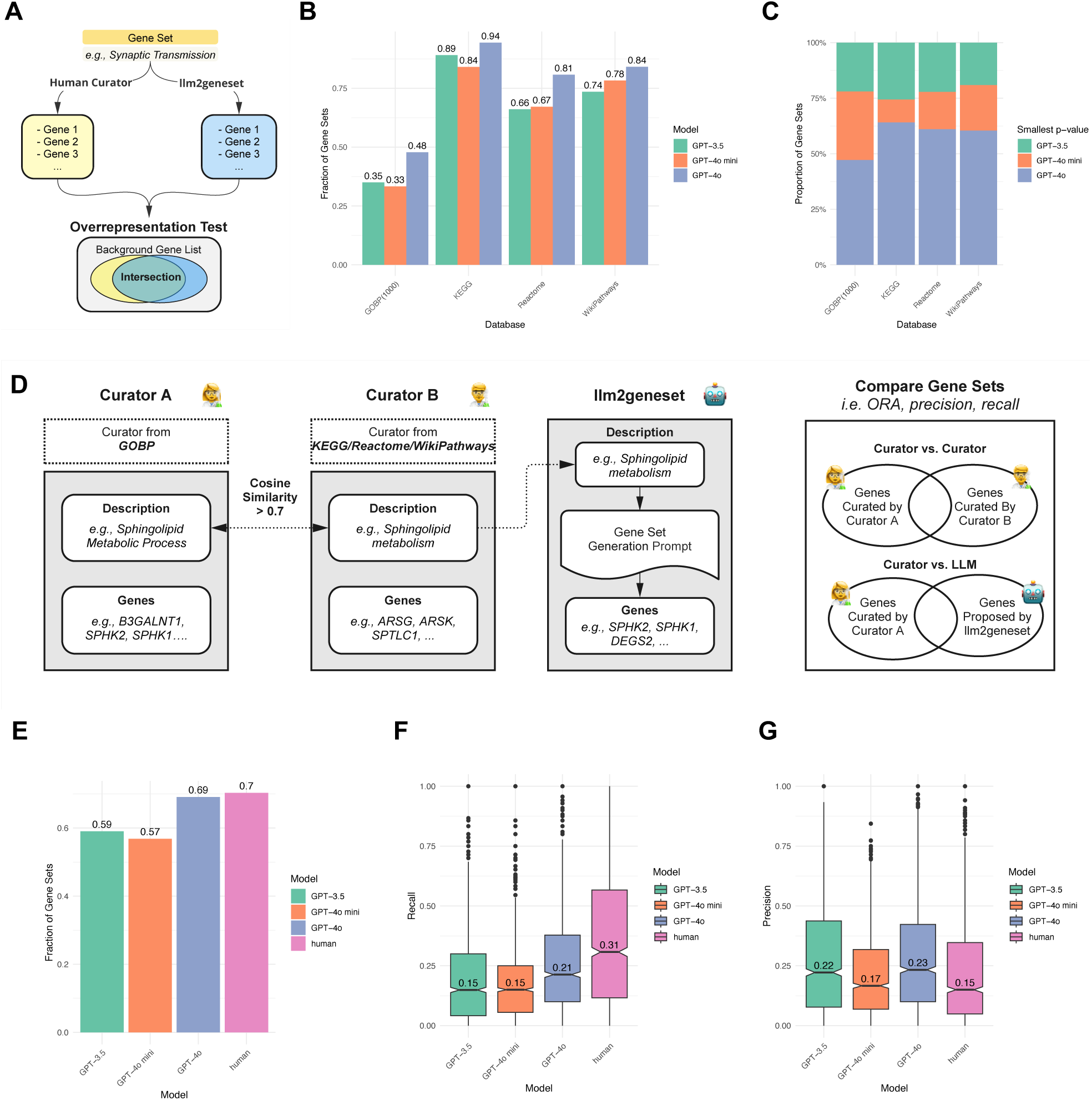
Evaluating gene sets generated by LLMs. (a) Gene sets generated by LLMs were assessed for significant overrepresentation in human-curated gene sets. Significant overrepresentation indicates that the LLM accurately associates gene set descriptions with corresponding genes. (b) Fraction of gene sets showing significant enrichment in curated sets at a Bonferroni-adjusted p-value of 0.01 across databases and LLMs. Here GOBP(1000) designates 1000 randomly sampled gene sets from GOBP. (c) Proportion of gene sets where the smallest overrepresentation p-value is associated with a specific model. (d) We identified 1,418 gene set descriptions with cosine similarity > 0.7 between the Gene Ontology Biological Process (GOBP) database and KEGG, Reactome, and WikiPathways. These gene sets allowed comparisons between human curators and established a baseline for human performance in gene set annotation. Gene sets were generated using LLMs based on KEGG, Reactome, and WikiPathways descriptions. (e) Fraction of gene sets significantly enriched in the GOBP reference across LLMs and human curators. “Human” refers to curator B from panel (d). (f) C (g) Boxplots showing recall and precision of gene sets generated by human curator B and LLMs.

We next sought to compare how LLMs might perform head-to-head with human curators. To do so, we focused on 1,418 gene sets in KEGG, Reactome, and WikiPathways (query gene sets) that had highly similar descriptions (cosine similarity > 0.7 using a text embedding model, see Methods) to the gene set descriptions in the entire GOBP gene set database (reference gene sets) (see Fig. 2D). Although they have highly similar descriptions, they originate from different databases and thus could be assumed to have been curated by different individuals. We found that 70% of these query gene sets from KEGG, Reactome, and WikiPathways were significantly enriched in the GOBP reference at a Bonferroni adjusted p-value of 0.01 (Fig. 2E). GPT-4o performed similarly at this task, with 69% of gene sets enriched in the human GOBP reference. We also computed the precision and recall of genes across these gene sets for LLMs as well as humans (Fig. 2F-G). We found that GPT-4o generated gene sets more conservatively than human curators with lower recall (p < 10^-13^, Wilcoxon rank sum), but higher precision (p < 10^-12^, Wilcoxon rank sum).

The prompt we evaluated above relied on two additional components: a role prompt and a format specification to generate outputs that can be used for downstream applications. Role prompting is a strategy by which to shape the output of an LLM by asking it to take on a specific role (see Methods) [12]. We evaluated whether the role-prompt had any impact on the quality of the results. We found that the role-prompt had little impact on the evidence for overrepresentation of curated genes in LLM generated gene sets (Fig. S2C). Formatting is crucial for subsequent use of the gene sets generated by LLMs. We assessed how often each LLM used HGNC symbols, as directed in the prompt, by comparison to gene symbols in Ensembl (GRch38.p14, release 112). We found that on average 95% of the gene symbols returned by GPT-4o were HGNC gene symbols (Fig. S2D). This number was 88% for GPT-3.5 and GPT-4o mini. We additionally noticed a tendency of the models to return duplicate gene symbols for a given gene set description. We found that GPT-3.5 generated gene sets with one or more duplicate gene symbols across 13-24% of gene sets, which was greatly reduced for GPT-4o (5-6%) (Fig. S2E). Overall, the LLMs we evaluated followed formatting instructions enabling generation of HGNC gene symbols for downstream use.

Token use is a crucial metric for LLMs and directly tied to compute cost [19]. Input tokens and output tokens are considered differently given that input tokens can be cached to reduce compute cost [20]. We quantified the number of input and output tokens used to generate LLM version of each database and model (Fig. S2F). Given that GPT-4o and GPT-4o mini both use the same tokenizer, 707,058 tokens were used as input to generate the gene sets using the descriptions from KEGG, Reactome, and Wikipathways. We found that GPT-4o used the most tokens in its output across databases. 1,004,497 output tokens were required to generate 3,939 gene sets using GPT-4o. In contrast, GPT-4o mini required 756,7728 output tokens. Quantification of token usage indicates that the compute cost associated with gene set generation was minimal overall.

### Ensembling and Confidence Improve the Precision of Gene Set Generation

LLM prompting is an active area of research [12]. On question-and-answer (QCA) benchmarking data sets various prompting strategies have been explored to increase LLM performance. We evaluated 3 prompting strategies on the problem of gene set generation: model reasoning, model confidence, and ensembling (Fig. 3A, see Methods). Encouraging an LLM to provide reasoning for why and how it arrived at a particular answer has been demonstrated to be a powerful approach to improve performance on QCA datasets [21]. We modified our prompt to require that the model provide a single sentence for its rationale for why a gene was included in a gene set (see Methods). We additionally considered whether we could elicit the model’s confidence in whether it thought a gene belonged to a gene set. We required the model to indicate low, medium, and high confidence for each gene, limiting gene sets to high confidence genes. We also considered ensembling. We used multiple generations of the model with differing random seed values. Across generations, we identified genes that the model consistently associated with a given gene set description. When a gene is consistently provided as output for a query for a gene set description, this may represent fixed and highly confident knowledge that the LLM captures.

**Figure 3.**
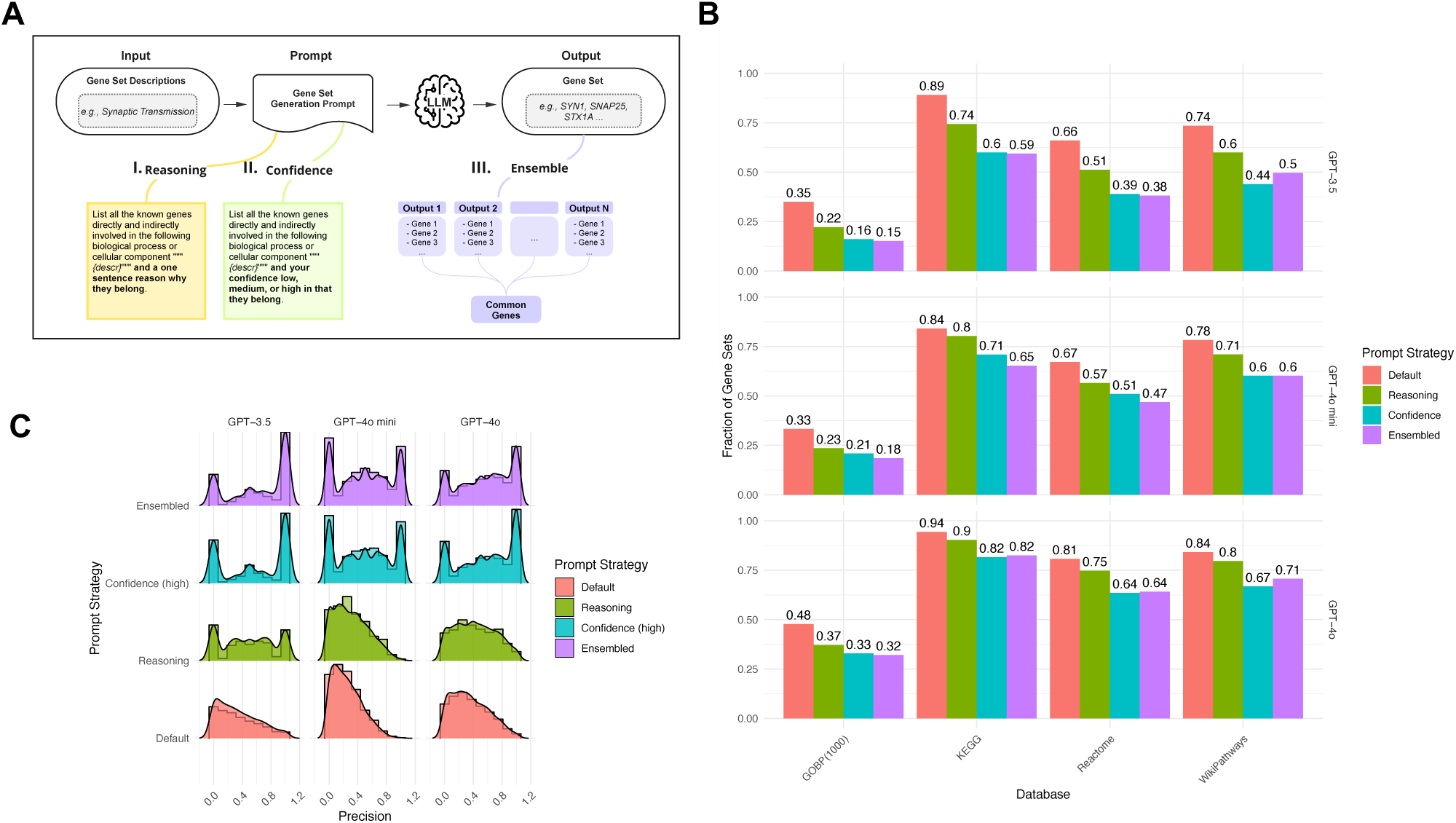
Assessment of Model Reasoning, Confidence, and Ensembling for Gene Set Generation. (a) Strategies to generate gene sets using an LLM prompt. (b) Fraction of genes where the LLM generated gene set is significantly overrepresented in the human curated gene set with the same natural language description. Bar plot color designates one of three prompting strategies illustrated in (a). (c) Precision of the gene sets returned or the fraction of LLM generated genes in a gene set that are in the human curated gene sets with the same description.

For each of these approaches reasoning, confidence, and ensembling, we observed a small decrease in the number of LLM generated gene sets that were overrepresented in human curated gene sets (Fig. 3B). The impact was more modest for GPT-4o. As expected, all of the gene sets generated by ensembling were the same size or smaller than those generated by our original prompt. We hypothesized that genes that appeared consistently in model generations with different seeds reflected the model’s higher confidence that the gene belonged to a given gene set. To quantify this, we evaluated whether these gene sets were more precise. Precision can be quantified by the number of genes in the LLM generated gene set belong to a curated gene set over the total number of genes returned. We found that both ensembling and model confidence, but not model reasoning increased the number of gene sets with high precision (Fig. 3C). These gene sets were only present when gene sets were restricted solely to high confidence genes, but not medium or low confidence genes (Fig. S3A). We examined the overlap between genes generated by ensembling and model confidence. Among gene sets that were significantly enriched in curated gene sets, we found a range of overlap between the gene sets generated by these approaches indicating that ensembling yields different genes than model confidence (Fig S3B).

Each of the strategies we considered requires additional tokens, thus increasing cost. As expected, token usage was highest for the ensembling strategy followed by model reasoning (Fig. S3C-S3D). Taken together, our results illustrate that prompting strategies can tune the precision of gene sets generated by LLMs.

### LLM Generated Gene Sets Enable Discovery of Multiple Enriched Biological Processes

LLMs have recently been proposed as tools to discover biological processes represented by an input set of genes [9]. In this setting, an LLM prompt includes a set of genes and asks an LLM to generate natural language description of the biological process these genes participate in – the reverse of our evaluation approach above. The prompt is evaluated based on how closely the LLM can recover the ground truth natural language description of the gene set in the output. In contrast to simple LLM prompting, *llm2geneset* generates a custom gene set database given a query gene set (Fig. 1). As LLMs may not use the same exact description as a curator used for a gene set, agreement can be quantified by, for example, how many words (unigrams) or word pairs (bigrams) from the ground truth description are present in the output of the LLM gene description.

Agreement can also be quantified by use of text embedding models and computation of cosine similarity between an embedding of the LLM description and an embedding of the ground truth description for a gene set [22].

We initially established a baseline using traditional overrepresentation analysis (ORA). To do so, we used as query gene sets from KEGG, Reactome, and WikiPathways. We then used the GOBP database to identify (from GOBP) gene sets that were significantly enriched in the query gene sets (from KEGG, Reactome, and WikiPathways). We compared the “ground truth” descriptions of query gene sets (from KEGG, Reactome, and Wikipathways) to the top-5 significant gene sets returned by performing ORA using GOBP (Fig. 4A). Across these significant gene set descriptions from GOBP, we used the maximum shared unigrams, bigrams, and cosine similarity between the “ground truth” descriptions from the original query gene sets to assess the quality of the results.

**Figure 4.**
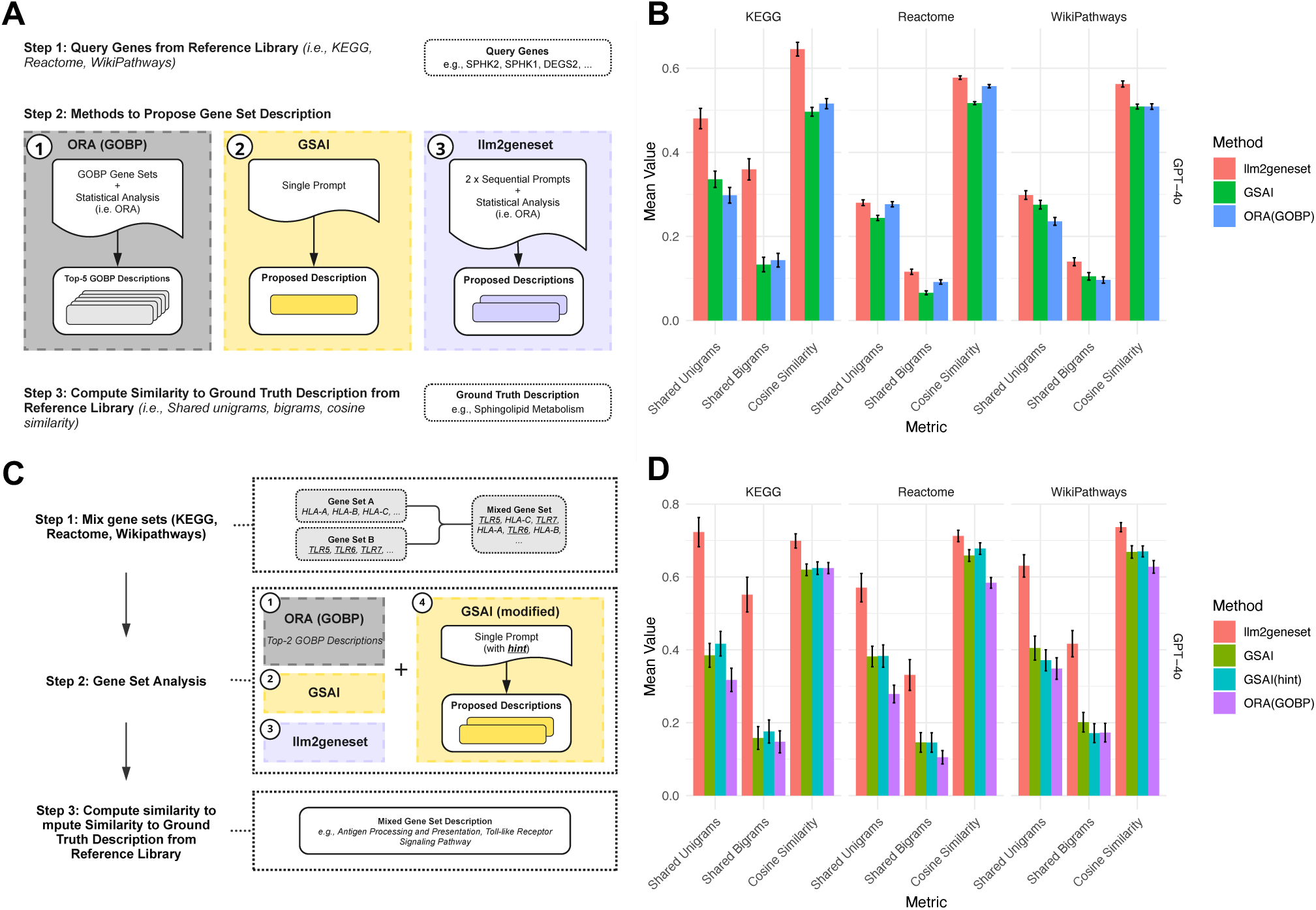
Gene Set Proposal Enables Discovery of Multiple Enriched Biological Functions. (a) Evaluation approach for traditional ORA using the GOBP database, GSAI, and *llm2geneset*. An input set of genes was provided. Each method was then allowed to propose gene set descriptions. These descriptions were compared using shared unigrams, bigrams, and cosine similarity with the ground truth description. (b) Mean fraction of shared unigrams (words) and bigrams (word pairs) and mean cosine similarity between gene set descriptions returned by the GSAI prompt and *llm2geneset* across databases for the GPT-4o model. Error bars designate the standard error of the mean. (c) To evaluate the ability of the approaches to identify multiple gene functions, we mixed gene sets that were easily identified by the approach in (a) and asked if each method recovered the mixed gene set descriptions. We also tested a modified GSAI prompt where we provided a hint indicating that 2 gene sets were present. (d) Mean fraction of shared unigrams (words) and bigrams (word pairs) and mean cosine similarity between gene set descriptions returned by GSAI, traditional ORA using GOBP, *llm2geneset* and the gene set descriptions originally assigned to mixed gene sets. Error bars designate the standard error of the mean.

We also compared our approach to an LLM prompting strategy called GSAI[9]. In contrast to our approach, GSAI used a single prompt to request the LLM return the biological process the input genes belong to along with both reasoning and confidence. Using benchmarking gene sets from KEGG, WikiPathways, and Reactome, we found that *llm2geneset* generated gene set descriptions had on average a higher fraction of shared unigrams and bigrams as well as higher cosine similarity with the ground truth gene set descriptions than GSAI and outperformed the ORA baseline (Fig. 4B, S4A). This observation was highly significant for the GPT-4o model using shared bigrams (Wilcoxon rank sum, p < 10^-13^ for KEGG, p < 10^-5^ for Reactome, and p < 10^-5^ for WikiPathways) and cosine similarity (Wilcoxon rank sum, p < 10^-11^ for KEGG, p < 10^-17^ for Reactome, and p < 10^-9^ for WikiPathways) as evaluation metrics. In terms of computational cost, our approach required a comparable number of input tokens to GSAI, but a larger number of output tokens (Fig. S4B).

Experiments often influence multiple biological processes, leading to the discovery of distinct overrepresented gene sets that include DEGs. To simulate such a scenario, we selected gene sets where the biological processes were recovered by both the GSAI prompt and *llm2geneset* with high similarity to the ground truth gene set description (> 0.7 cosine similarity) for each benchmarking gene set database. Next, we combined 50 pairs of these gene sets from each gene set database. We then evaluated the recovery of the distinct biological processes from these mixed gene sets, comparing the GSAI prompt with our approach across LLMs (see Methods).

We established a baseline using traditional ORA by measuring text similarity between of the concatenated text of the top-2 significant gene sets returned from the GOBP database. We found that with the GSAI prompt LLMs were frequently unable to identify multiple gene functions in a gene set. This also held when we modified the original GSAI prompt to include a text “hint” that indicated that two distinct gene sets were present in the input query gene set (see Methods).

Compared to both GSAI, GSAI with a two gene set hint, and the ORA baseline, *llm2geneset* more frequently recovered multiple functions as reflected by higher fraction of shared unigrams and bigrams as well as cosine similarity (Fig. 4D, Fig. S4C). This observation was highly significant for the GPT-4o model using shared bigrams (Wilcoxon rank sum, p < 10^-11^ for KEGG, p < 10^-4^ for Reactome, and p < 10^-4^ for WikiPathways) and cosine similarity (Wilcoxon rank sum, p < 10^-3^ for KEGG, p < 10^-5^ for Reactome, and p < 10^-5^ for WikiPathways) as evaluation metrics. Our analysis indicates that coupling generation of gene sets by LLM to overrepresentation analysis is a more effective strategy than model prompting to uncover multiple distinct gene set functions within a set of genes.

### LLM Based Gene Set Generation Enables Steerable Overrepresentation Analysis

After establishing that our LLM-based approach is more effective than using a single prompt to identify multiple biological processes within a gene set, we applied this method to the differentially expressed genes (DEGs) from an experiment involving microglia differentiated from induced pluripotent stem cells (iMGs), treated with AL002 (N=4) compared to an isotype control (N=4) (see Methods). AL002 is an investigational, humanized, TREM2-selective, agonistic monoclonal antibody in Phase 2 trials for treatment of early Alzheimer’s disease [23]. Activation of microglia is hypothesized to be an important mechanism of AL002. In iMGs, we found that 30 genes were differentially expressed at an FDR of 10% between iMG that received AL002 to the isotype control (Fig. 5A).

**Figure 5.**
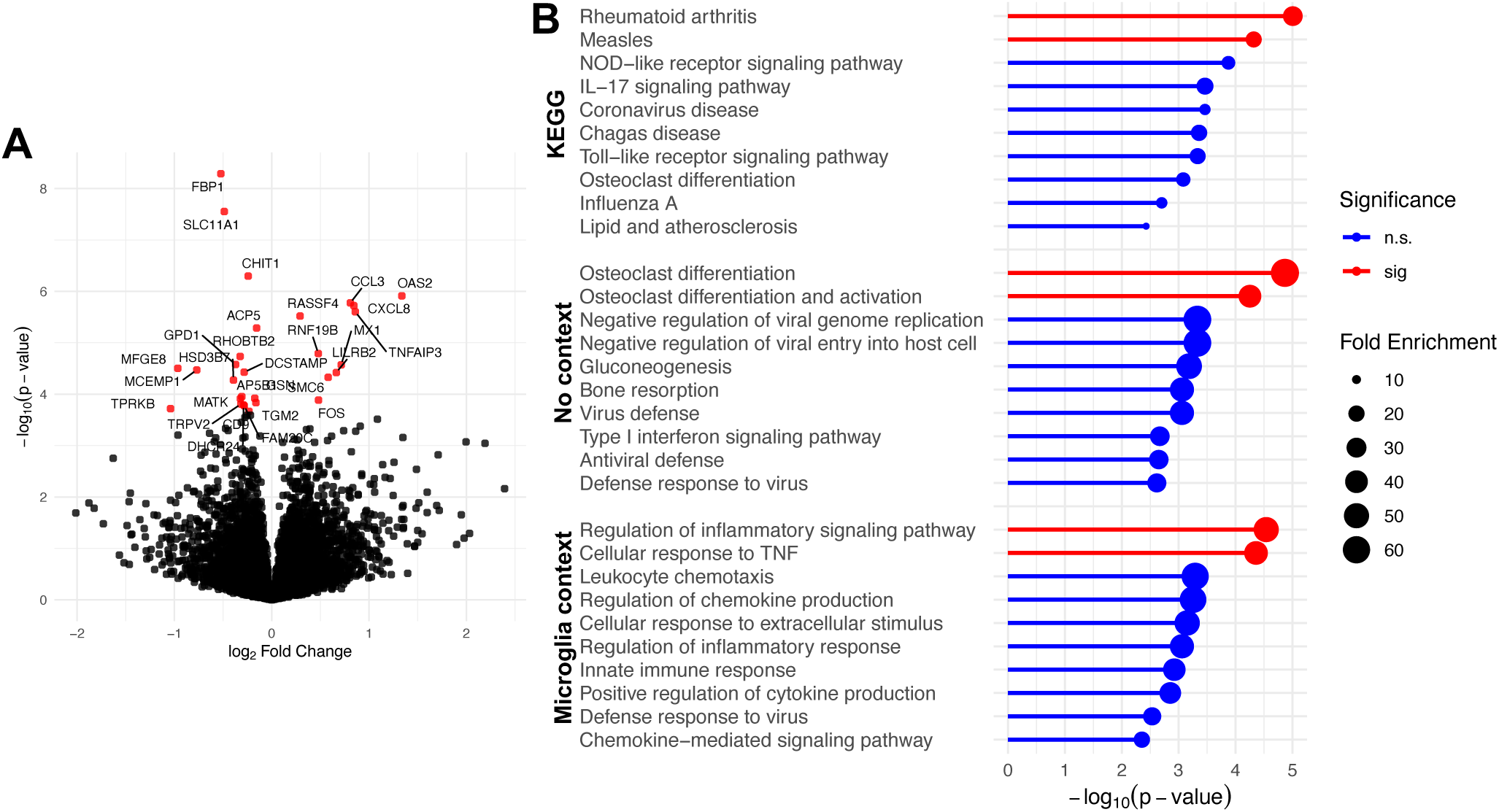
Comparison of overrepresentation analyses using static and LLM generated gene sets. (a) Volcano plot of DEGs comparing iPSC derived microglia treated with a TREM2 agonist antibody (AL002) to an IgG control antibody at 1ug/ml for 24 hours. Differentially expressed genes at FDR 10% are shown as red points. (b) Overrepresentation analysis results for biological process and pathways in: (top) the KEGG database; (middle) GPT-4o generated gene set database using no contextual information about the experiment; (bottom) GPT-4o generated gene set database using the contextual text “in vitro microglia treated with a TREM2 agonist antibody.” The lollipop plot indicates the -log10(p-value) for overrepresentation of the gene sets in the set of DEGs in (a). Size of the circles in the plot designates fold enrichment of the gene set. Significant (sig) gene sets at an FDR of 1% are shown in red. Non-significant (n.s.) gene sets are shown in blue.

Given that it is difficult to define ground truth overrepresented gene sets from RNA-seq data, we qualitatively compared the overrepresentation results of using a fixed database derived from KEGG to that of results of *llm2geneset* (Fig. 5B). Using the KEGG database, we found several highly ranked pathways that were irrelevant to microglia. These include pathways rheumatoid arthritis and measles which provide little information on the impact of AL002 *in vitro*. In contrast, a larger number of highly ranked pathways related to immune and osteoclast function were detected by *llm2geneset*. The enrichment in osteoclast function was driven by DEGs such as *ACP5* and *DCSTAMP*, which have also been associated with functions in the human immune system [24–26]. In this setting, the LLM was unaware of the experimental context of the input DEGs.

To address this scenario, we leveraged the ability of *llm2geneset* to steer the model toward gene set descriptions related immune and microglial function (see Methods, Fig. 5B). Here, the gene set proposal prompt included the context “in vitro microglia treated with a TREM2 agonist antibody.” Using this approach we found the biological process “regulation of inflammatory responses” was significant at FDR 1%. This result is consistent with TREM2’s function of the innate immune system in the brain [27]. Pathways and biological processes related to immune function were ranked highly overall and gene sets related to osteoclast function were not tested. Overall, this example illustrates that LLMs can be steered to generate gene set databases highly relevant to a specific RNA-seq experiment. These gene sets can be evaluated for overrepresentation in a set of DEGs to facilitate interpretation of RNA-seq data sets and subsequent hypothesis generation.

## Discussion

The ability to dynamically generate gene set databases using LLMs, tailored to input genes and contextual information, offers distinct advantages for gene set enrichment analysis. This approach enables researchers to customize gene sets to the unique characteristics of their data and scientific questions, addressing a key limitation of static, pre-defined databases. Moreover, because LLMs are continuously trained on the evolving scientific literature, they can adaptively update these gene sets as scientific knowledge progresses, providing a potential solution to the challenges associated with maintaining an up-to-date gene set database [28]. Additionally, *llm2geneset* can be seamlessly integrated into existing gene set enrichment analysis methods, such as GSEA, Enrichr, or CAMERA. The generated databases can be treated as static by these methods, enabling them to leverage the flexibility and adaptability of our framework while maintaining compatibility with established approaches. To broaden accessibility, a web application (https://llm2geneset.streamlit.app), enables researchers to use *llm2geneset* without programming expertise, expanding its utility for experimental biologists.

LLMs have been demonstrated to possess emerging planning and reasoning capabilities [29]. A single prompt, given a set of input query genes such as DEGs, can leverage these reasoning capabilities to infer the function of the input genes. However, we find that this approach is less effective than *llm2geneset* at inferring the function of a given input set of genes, particularly when multiple gene sets are mixed. This advantage arises from our framework’s ability to propose and evaluate multiple candidate gene sets against the input query, enabling the discovery of distinct biological functions within complex gene sets. Additionally, a single prompt is more susceptible to being influenced by researchers steering the model too strongly through prompt phrasing. In contrast, steering within *llm2geneset* impacts only the proposed gene set descriptions, not the gene sets themselves, as the framework validates all generated gene sets against the input query genes.

Our work has several limitations. Certain gene sets cannot be recovered using LLM prompting alone. Integrating retrieval-augmented generation may enhance performance, but careful benchmarking is necessary to validate improvements[30]. Our comparison between LLMs to human curators is limited to 1,418 gene sets. How human curators perform relative to LLMs needs further study. Additionally, our study evaluates only three large language models - GPT-3.5, GPT-4o mini, and GPT-4o - and does not assess reasoning models on the tasks defined here[31].

Differences in model architectures, training approaches, and inference algorithms processes must be further examined using the gene set tasks and ORA baselines introduced in this work. Some gene sets are molecularly defined based on transcriptomics data, and our direct prompting approach may not recover these sets[32]. However, LLMs’ ability to generate code could enable automated creation of such gene sets, or models trained on expression data may address this limitation[33,34]. Despite these challenges, we anticipate LLMs will be essential tools for deriving insights from high-dimensional transcriptomics and proteomics data sets.

## Methods

### LLMs and Gene Sets Databases

We used the following versions of OpenAI large language models: GPT-4o mini (gpt-4o-mini-2024-07-18), GPT-3.5 (gpt-3.5-turbo-0125), and GPT-4o (gpt-4o-2024-08-06). GMT files containing gene sets were downloaded from https://maayanlab.cloud/Enrichr/#libraries. We used the Reactome (Reactome_2022), KEGG (KEGG_2021_Human), WikiPathways (WikiPathway_2023_Human), and the Gene Ontology Biological Process (GOBP, GO_Biological_Process_2023) gene set databases for evaluation. Each of the gene set descriptions were cleaned to remove any unique identifiers. To favor reproducible outputs from the LLMs, we used the seed parameter in the model API. We obtained the list of valid human HGNC symbols using the biomaRt R package by querying the `hsapiens_gene_ensembl` database for protein coding gene symbols [35].

### Gene Set Generation

We used the following prompt to generate gene sets:

`List all the known genes directly and indirectly involved in the following biological process or cellular component “““{descr}”””. Use the following JSON schema:`

~~~
‘‘‘json
{{
     “type”: “array”,
     “items”: {{
       “type”: “object”,
       “properties”: {{
           “gene”: {{
               “type”: “string”,
           }}
      }},
      “required”: [“gene”]
    }}
}}
‘‘‘
The field ‘genè is a gene involved in the following biological process or cellular component: “““{descr}”””. Use the HUGO Gene Nomenclature Committee (HGNC) gene abbreviations. Place the output in a JSON code block. Do not add any comments in the JSON code block.
~~~

In the above prompt `{descr}` was replaced with a natural language description of a gene set. We used the following role prompt which we placed in the system message:

`You are an expert in cellular and molecular biology.`

The https://pypi.org/project/json-repair/ package was used to parse out the returned genes and repair any minor JSON formatting errors. If the JSON output was not repairable, we queried the model again with a different seed value. We used the input and output tokens returned by the model API, accumulating additional tokens if additional model queries were needed. We removed any duplicate gene symbols returned by the LLM. The above prompt, query to an LLM, and parsing can be concisely written as function that takes as input a string description of a gene set and returns a string list with the corresponding gene symbols:

`G:List[str] <-GetGenes(D:str)`

### Evaluating Gene Sets Generated by LLM

We evaluated the gene sets generated using the prompt above by examining the overrepresentation gene sets generated by LLM in the gene set curated by a human with the same description. We computed the overlap between the gene sets returned by LLM and the human curated gene sets and used the hypergeometric distribution to perform a one-tailed Fisher’s exact test to compute a p-value corresponding to the probability whether the overlap, or higher overlap, was observed by chance. We used the hypergeom.sf in the scipy.stats package to compute numerically accurate p-values. We assumed possible set of N=19,846 human protein coding genes based on the statistics of GRCh38.p14 (https://useast.ensembl.org/Homo_sapiens/Info/Annotation). We corrected for multiple testing across gene sets using Bonferroni correction or false discovery rate as indicated. To evaluate model generations, we report the fraction of significantly overrepresented gene sets after multiple testing correction. Multiple testing corrections were performed across all gene sets within a database.

### Comparison of LLM and Human Curator Performance

To evaluate the performance of large language models (LLMs) relative to human curators, we focused on 1,418 gene sets from KEGG, Reactome, and WikiPathways (query gene sets) with highly similar descriptions to gene sets in the GOBP database (reference gene sets). Similarity was determined using cosine similarity (>0.7) computed using OpenAI’s text-embedding-3-large model. These query gene sets and reference gene sets were assumed to be independently curated as they originate from different databases. For each query gene set, we performed overrepresentation analysis (ORA) against the GOBP database, considering a Bonferroni-adjusted p-value threshold of 0.01 for significance. GPT-4o was evaluated using the same approach to generate gene sets for comparison. Additionally, precision and recall were computed for the genes included in the gene sets generated by LLMs and human curators. Statistical differences between LLMs and human curators were assessed using the Wilcoxon rank-sum test.

### Additional Gene Set Generation Prompting Strategies

Prompts for confidence and reasoning closely followed our original prompt above. We modified the query portion of the original prompt to elicit model confidence as follows:

`List all the known genes directly and indirectly involved in the following biological process or cellular component “““{descr}””” and your confidence low, medium, or high in that they belong.`

The formatting portion of the prompt included modifications to the output JSON schema to capture this additional confidence information. Only high confidence genes were used in our evaluation. Model reasoning was elicited in a similar manner with the query portion of the prompt modified as follows:

`List all the known genes directly and indirectly involved in the following biological process or cellular component “““{descr}””” and a one sentence reason why they belong.`

Here, we also modified the JSON schema in the formatting portion of the prompt to output such that the single sentence corresponding to the model reason was captured. Ensembling was conducted by performing 5 generations of gene sets with different seed values for the model to obtain different genes for each generation. Genes that were consistently observed in all 5 generations were returned as the genes for the given gene set description.

### Proposing Gene Set Descriptions from a Set of Genes

We used the following prompt to propose a set of gene set descriptions from a list of genes.

List {n_pathways} biological pathways, biological processes, or cellular components that contain the following genes “““{genes}””” with high confidence. Be as specific as possible. List non-overlapping pathways, processes, or components. Do not include the gene names in the outputs. Use the following JSON schema:

~~~
‘‘‘json
{{
     “type”: “array”,
     “items”: {{
        “type”: “object”,
        “properties”: {{
            “p”: {{
                “type”: “string”,
            }},
      }},
      “required”: [“p”]
    }}
}}
‘‘‘
Example output will look like the following:
‘‘‘json
[{{“p”:“BP or Pathway 1”}},
{{“p”:“BP or Pathway 2”}},
{{“p”:“BP or Pathway 3”}},
{{“p”:“BP or Pathway 4”}}
‘‘‘
~~~

The element ‘p’ designates a pathway, biological process or cellular component. Place the output in a JSON code block. Do not add any comments in the JSON code block.

In the above prompt, `{n_pathways}` was replaced with the number of desired biological pathways and process descriptions. `{genes}` was replaced by a comma separated list of genes for which we requested these biological processes and pathways. As with the gene set generation prompt, we used the https://pypi.org/project/json-repair/ package to parse out the returned genes and repair any minor JSON formatting errors. If the JSON output was not repairable, we queried the model again with a different seed value.

To steer the model toward experimentally relevant biological processes and pathways we additionally modified the above prompt to include contextual information when this contextual information was provided:

List {n_pathways} biological pathways, biological processes, or cellular components that contain the following genes “““{genes}””” with high confidence. Also consider the following context as related to the genes: “““{context}””” when selecting pathways, processes, and components.

Here `{context}` was replaced with a user provided string that provided additional context to steer the model generations (e.g. “in vitro microglia treated with a TREM2 agonist antibody”). The above can be concisely written as a function that takes as input a list of genes D, a requested number of processes and pathways N, and optionally a C context string:

P:List[str] <-GetPathwaysProcesses(D:List[str], N:int, C:str)

### Discovery of Multiple Enriched Biological Processes in Gene Sets

*llm2geneset* proposes pathways and biological process descriptions based on the input set of genes and an experimental context. These pathway descriptions, alone, are used to generate gene sets which are then tested for overrepresentation in the input set of DEGs. The `GetGenes()` function call below does not have access to any previous context. The parameters of the algorithm are as follows:

- D = set of DEGs (or any set of genes)
- N = number of biological pathways and processes to propose
- C = (optional) contextual information regarding the experiment from which the DEGs were obtained
- B = number of background gene sets for overrepresentation analysis Pseudo-code is provided below:

llm2geneset(D:List[str], N:int, C:str) R = []

P = GetPathwaysProcesses(D) for pathway in P:

G = GetGenes(pathway)

p = hgsf(|intersect(D,P)|-1, B, |G|, |D|) R = R.append((pathway,p))

return padjust(R)

In the above pseudo-code, the function `padjust()` computes q-values to account for multiple testing. `hgsf()` is one minus the cumulative distribution function (1-CDF) of the hypergeometric distribution. The function call computes the p-value according to a one-tailed Fisher’s exact test. *llm2geneset* returns a list of N pathways sorted on their overrepresentation p-values.

### GSAI Prompt Details

We used the previously published GSAI prompt [9]. The GSAI prompt provides extensive instructions to an LLM to provide a gene set description, a confidence value, and an analysis of how genes belong to the returned gene set description. We used regular expressions to parse out the gene set name, confidence, and the analysis. When we were unable to parse these outputs, we queried the LLM again with a different seed value to obtain outputs that could be parsed by regular expressions.

In our experiments with mixed gene sets below, we added the following “hint” to the GSAI prompt as the second line:

There are 2 distinct biological processes performed by this system of interacting proteins.

### Evaluating Biological Process Discovery from Gene Sets

We compared the performance of GSAI and *llm2geneset* through two experiments. In the first experiment, we used gene sets from KEGG, Reactome, and WikiPathways to evaluate how closely the outputs of the two methods matched the original gene set descriptions. The input gene sets were shuffled before being submitted to either GSAI or *llm2geneset*. We assessed similarity using three metrics: (1) the fraction of single words (unigrams) and (2) word pairs (bigrams) in the original descriptions that appeared in the returned descriptions, and (3) cosine similarity between text embeddings. Embeddings were computed using OpenAI’s text-embedding-3-large model, and the dot product of the embeddings was used for comparison. The average similarity score across all gene sets was used to evaluate the two methods. For *llm2geneset*, which generates multiple gene sets (N=5), we only considered gene sets significant at a Bonferroni-corrected p=0.002. If no significant sets were found, we returned “None Found.” The maximum score for each metric across the significant gene set descriptions was reported.

In the second experiment, we focused on gene sets where the cosine similarity between GSAI and *llm2geneset* descriptions was >0.7 in the first experiment. We randomly selected 50 such pairs, combined their genes, and shuffled the new gene sets. For GSAI, we used the parsed name from the LLM output, and for *llm2geneset*, we generated N=5 gene sets, again keeping only those significant at p=0.002. We combined the descriptions of the significant gene sets into a comma-separated string or returned “None Found” if no significant sets were identified. We compared the GSAI description and the *llm2geneset* description to a reference string, which combined the original human-curated descriptions in a comma-separated format. Similarity was quantified by the fraction of unigrams and bigrams from the reference string present in the GSAI and *llm2geneset* outputs, as well as the cosine similarity between the strings. The average similarity across all 50 combined gene sets was used to compare the two methods.

### Bulk RNA-seq from iMGs Treated with AL002

Induced pluripotent stem cells (iPSCs) were differentiated into mature microglia as previously described [36]. A total of 2.5 x 10^4^ cells each well were seeded in a 96-well plate format, and after 7 days in culture, the cells were treated with AL002 or an isotype-IgG control at 1ug/ml and incubated for an additional 24h. Next, cells were harvested, and RNA was isolated using the RNeasy 96 QIAcube HT Kit (Qiagen, Hilden, Germany). The quality of RNA was assessed using the TapeStation System (Agilent Technologies, Santa Clara, CA). RNA samples were converted to cDNA libraries using the Universal Plus mRNA-Seq kit with NuQuant (Tecan Genomics, Redwood City, CA). RNA sequencing was performed on an Illumina NovaSeq 6000 system using a NovaSeq V1.5 kit (SeqMatic, Fremont, CA), with 2×100 bp paired-end reads.

Sequencing quality was evaluated using FastQC (v0.12.1). Raw reads were trimmed and filtered using fastp (v0.23.4) to remove low quality bases and adapter sequences, discarding reads shorter than 25 bp[37]. Trimmed reads were aligned to the reference GRCh38 transcriptome with STAR (v2.7.11b)[38]. Sequencing and alignment quality control was summarized with MultiQC (v1.19)[39]. Differential expression analysis was conducted using DESeq2[40].

## Code and Data Availability

All code, notebooks, and benchmarking data sets are available here: https://github.com/Alector-BIO/llm2geneset.

The prompts used in this study are available for download here: https://github.com/Alector-BIO/llm2geneset/tree/main/src/llm2geneset/prompts

A streamlit web application incorporating this tool is available here: https://github.com/Alector-BIO/llm2geneset/tree/main/webapp.

An instance of the web application can also be accessed here: https://llm2geneset.streamlit.app.

## Conflict of Interest Statement

All authors are employees of Alector Inc.

## Author Contributions

ZK and JZ conceived of the idea. ZK, JZ, RYW, and AM contributed ideas and analyses. XW, RA, and JK contributed data. All authors read and approved the final manuscript.

## Supplementary Figures

**Figure S2.**
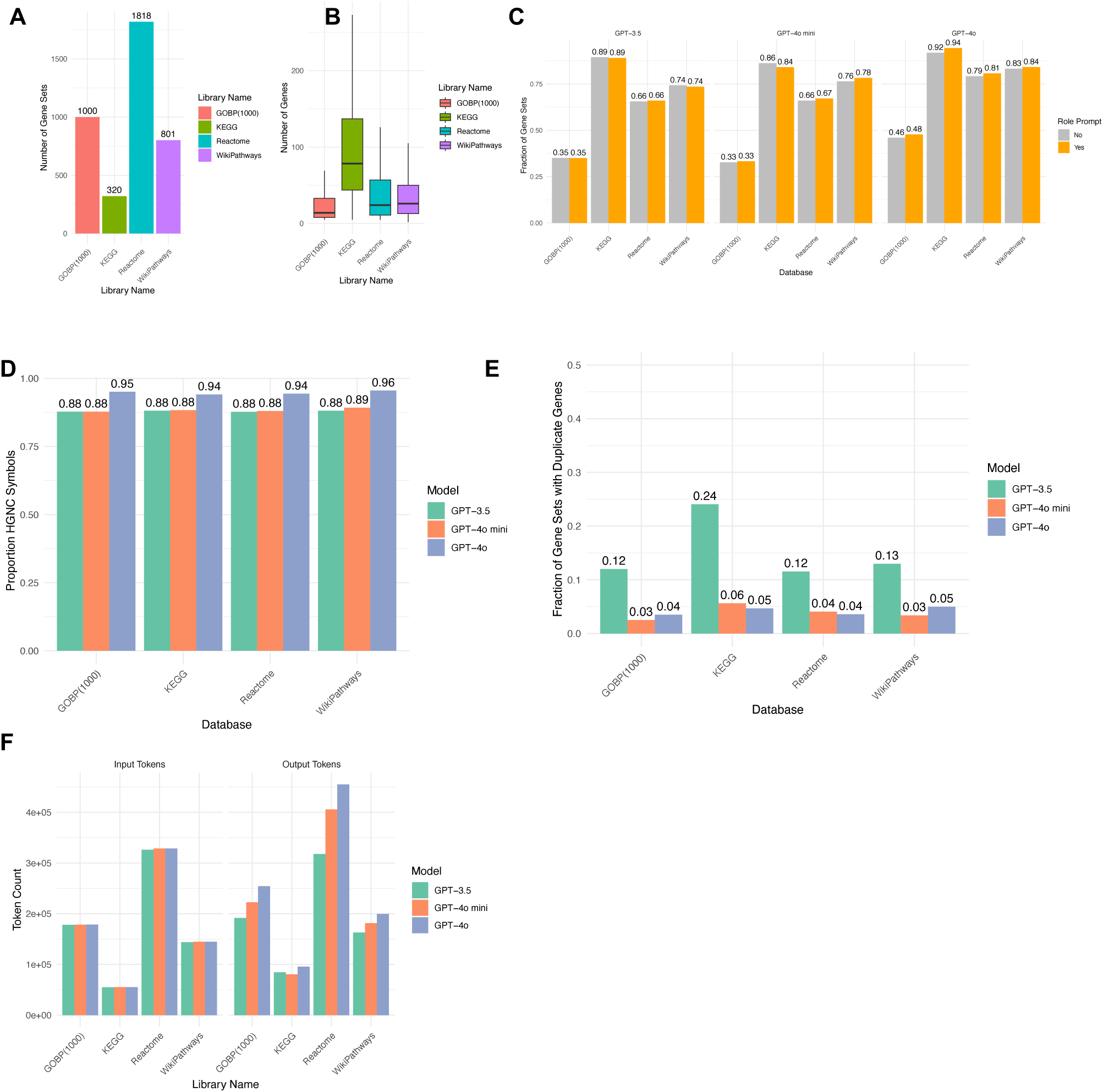
(a) Number of gene sets in the Reactome, WikiPathways and KEGG databases used for benchmarking. (b) Boxplot of the number of genes in each gene set across gene set databases. (c) Fraction of gene sets that show significant enrichment in human curated gene sets at a Bonferroni adjusted p-value of 0.01 across databases and LLMs. Plot compares the enrichment between gene sets generated using a role prompt and gene sets generated without a role prompt. (d) Fraction of gene symbols returned by LLMs across gene set databases that were known HGNC symbols in the Ensembl (GRch38.p14) genome annotation. (e) Fraction of gene sets that contain duplicate genes across LLMs for each of the benchmark gene sets. (f) Total number of input and output tokens used to generate gene sets across models and databases.

**Figure S3.**
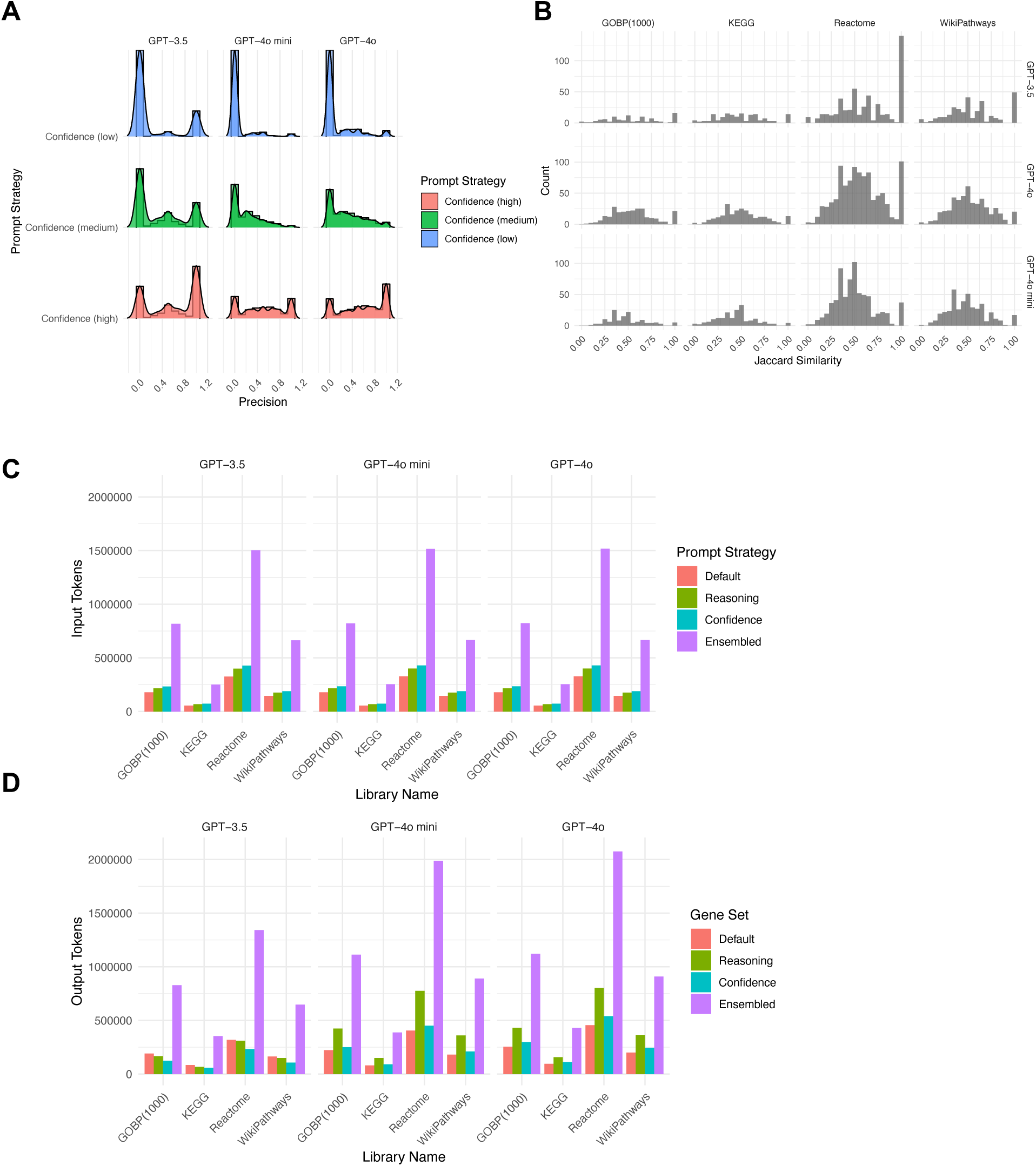
(a) Precision of gene sets comprising of only low, medium, and high confidence genes (b) Jaccard similarity between gene sets from the high confidence strategy and from ensembling that were significantly enriched in human curated gene sets at an adjusted p < 0.01 across databases and models. (c) Input tokens and (d) output tokens used by the default prompt strategy as compared to a prompting strategy that incorporates model reasoning, self-reported confidence, and ensembling.

**Figure S4.**
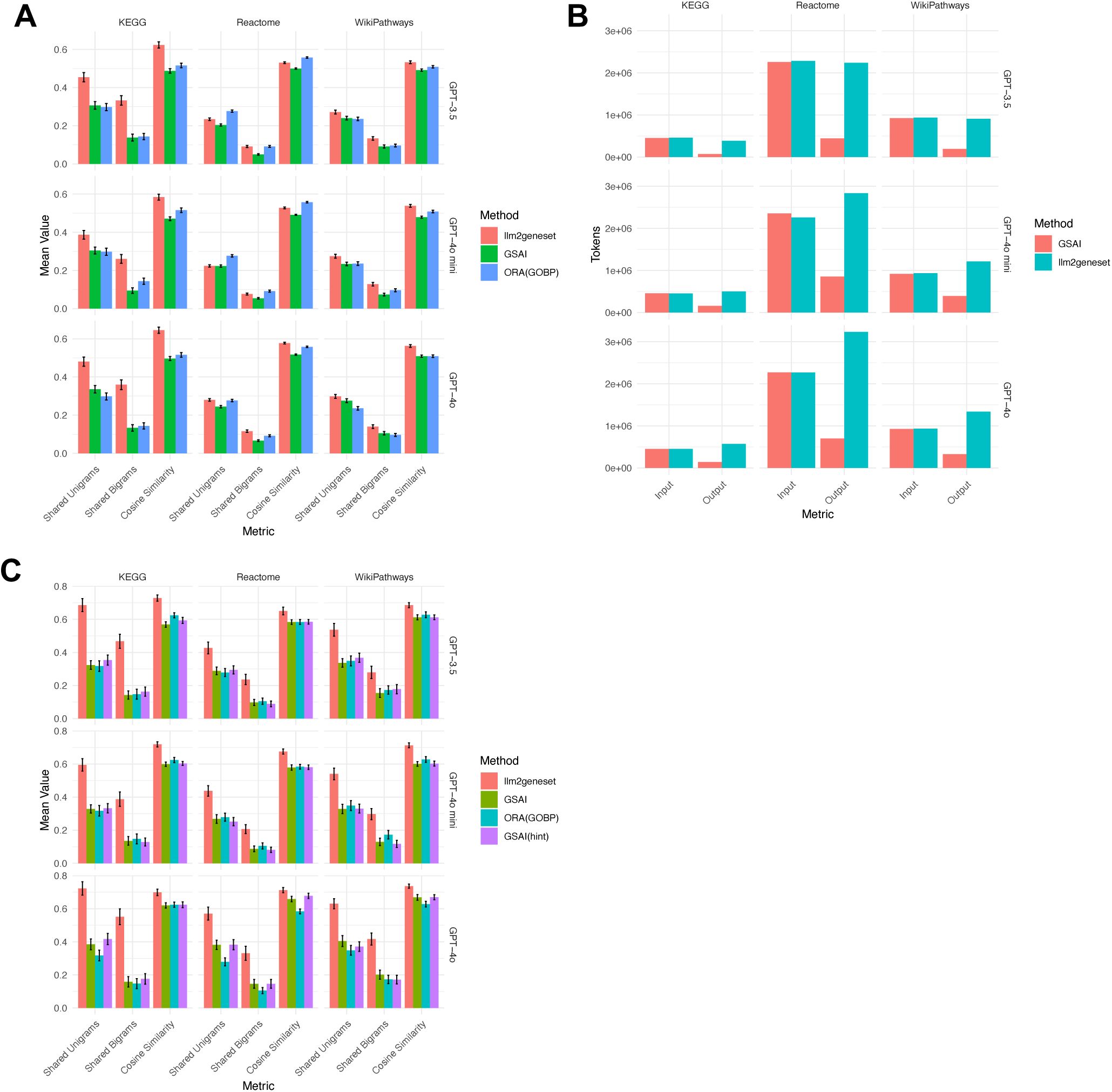
(a) Mean fraction of shared unigrams (words) and bigrams (word pairs) and mean cosine similarity between gene set descriptions returned by the GSAI prompt and *llm2geneset* across databases and LLMs. Error bars designate the standard error of the mean. (b) Input and output token usage of GSAI and *llm2geneset* across databases and models. (c) Mean fraction of shared unigrams (words) and bigrams (word pairs) and mean cosine similarity between gene set descriptions returned by GSAI and *llm2geneset* and the gene set descriptions originally assigned to mixed gene sets. Error bars designate the standard error of the mean.

## Notes

https://github.com/Alector-BIO/llm2geneset

### Competing Interest Statement

All authors are current employees of Alector, Inc.

### Summary of Updates

In this revision, we have included several additional experiments. First, we performed a head-to-head comparison of LLMs with human curators using gene sets from KEGG, Reactome, and WikiPathways that had highly similar descriptions (cosine similarity >0.7) to those in the Gene Ontology Biological Process (GOBP) database. This allowed us to evaluate LLM performance relative to human curation. Second, we established a baseline using traditional overrepresentation analysis (ORA) by comparing query gene sets (from KEGG, Reactome, and WikiPathways) to the top-5 significantly enriched GOBP gene sets. These results were compared against the ground truth descriptions of the query gene sets. Additionally, we revised the manuscript to improve readability, updated the introduction and discussion sections, and added figures to clarify the computational experiments.

